# The SARS-CoV-2 spike N-terminal domain engages 9-*O*-acetylated α2-8-linked sialic acids

**DOI:** 10.1101/2022.09.14.507904

**Authors:** Ilhan Tomris, Luca Unione, Linh Nguyen, Pouya Zaree, Kim M. Bouwman, Lin Liu, Zeshi Li, Jelle A. Fok, María Ríos Carrasco, Roosmarijn van der Woude, Anne L.M. Kimpel, Mirte W. Linthorst, Enrico C.J.M Verpalen, Tom G. Caniels, Rogier W. Sanders, Balthasar A. Heesters, Roland J. Pieters, Jesús Jiménez-Barbero, John S. Klassen, Geert-Jan Boons, Robert P. de Vries

## Abstract

SARS-CoV-2 viruses engage ACE2 as a functional receptor with their spike protein. The S1 domain of the spike protein contains a C-terminal receptor-binding domain (RBD) and an N-terminal domain (NTD). The NTD of other coronaviruses includes a glycan-binding cleft. However, for the SARS-CoV-2 NTD protein-glycan binding was only observed weakly for sialic acids with highly sensitive methods. Amino acid changes in the NTD of Variants of Concern (VoC) shows antigenic pressure, which can be an indication of NTD-mediated receptor binding. Trimeric NTD proteins of SARS-CoV-2, Alpha, Beta, Delta, and Omicron did not reveal a receptor binding capability. Unexpectedly, the SARS-CoV-2 Beta subvariant strain (501Y.V2-1) NTD binding to Vero E6 cells was sensitive to sialidase pretreatment. Glycan microarray analyses identified a putative 9-*O*-acetylated sialic acid as a ligand, which was confirmed by catch-and-release ESI-MS, STD-NMR analyses, and a graphene-based electrochemical sensor. The Beta (501Y.V2-1) variant attained an enhanced glycan binding modality in the NTD with specificity towards 9-*O*-acetylated structures, suggesting a dual-receptor functionality of the SARS-CoV-2 S1 domain, which was quickly selected against. These results indicate that SARS-CoV-2 can probe additional evolutionary space, allowing binding to glycan receptors on the surface of target cells.

**Graphical abstract:** 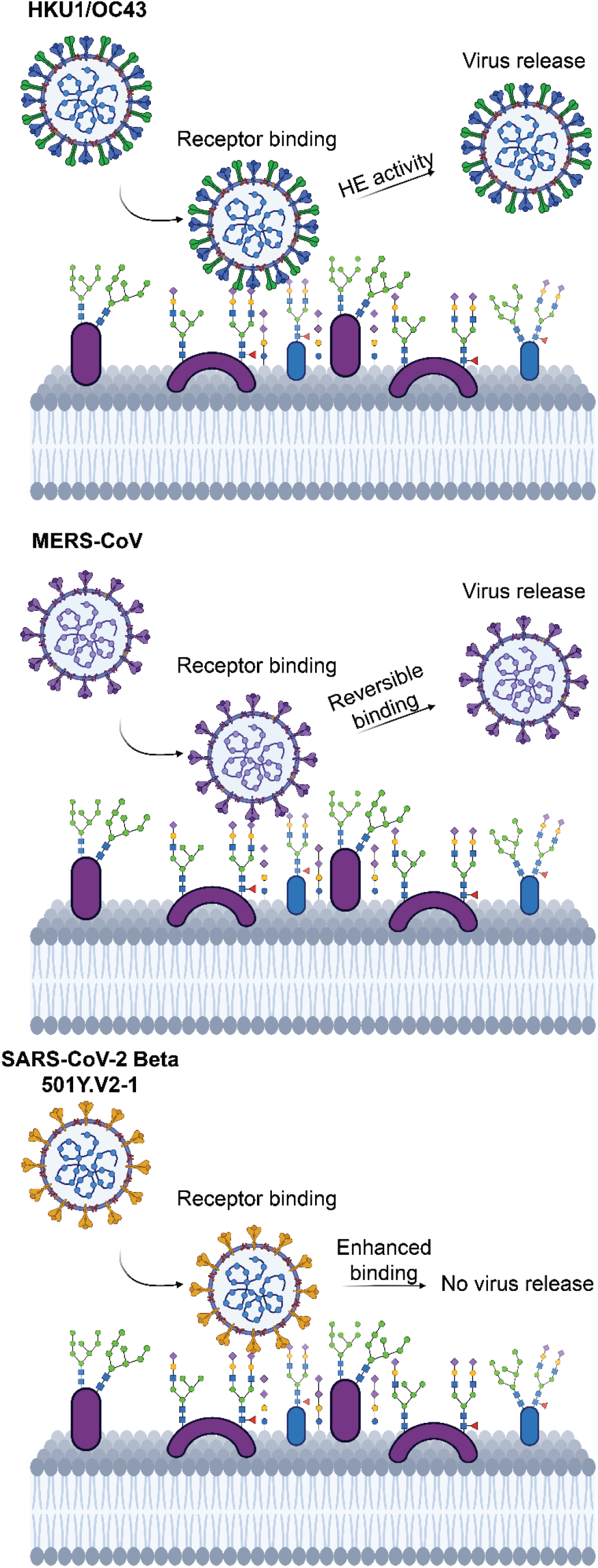

**Synopsis:** Coronaviruses utilize their N-terminal domain (NTD) for initial reversible low-affinity interaction to (sialylated) glycans. This initial low-affinity/high-avidity engagement enables viral surfing on the target membrane, potentially followed by a stronger secondary receptor interaction. Several coronaviruses, such as HKU1 and OC43, possess a hemagglutinin-esterase for viral release after sialic acid interaction, thus allowing viral dissemination. Other coronaviruses, such as MERS-CoV, do not possess a hemagglutinin-esterase, but interact reversibly to sialic acids allowing for viral surfing and dissemination. The early 501Y.V2-1 subvariant of the Beta SARS-CoV-2 Variant of Concern has attained a receptor-binding functionality towards 9-*O*-acetylated sialic acid using its NTD. This binding functionality was selected against rapidly, most likely due to poor dissemination. Ablation of sialic acid binding in more recent SARS-CoV-2 Variants of Concern suggests a fine balance of sialic acid interaction of SARS-CoV-2 is required for infection and/or transmission.

## Introduction

ACE2 is widely recognized as the functional and essential receptor of SARS-CoV-2 [1, 2]. Several reports also demonstrate the importance of the glycocalyx, a thick layer of glycans covering every eukaryotic cell and a known barrier for a plethora of pathogens [3]. Within this glycocalyx, heparan sulfate moieties have been shown to be an important attachment factor for a variety of viruses including coronaviruses [4, 5]. Another element of the glycocalyx, sialylated glycans, are like heparan sulfates negatively charged and are essential receptors for a wide variety of viruses, again including members of the coronaviruses [6–8].

Interaction of the SARS-CoV-2 spike glycoprotein towards (sialo)glycans has been confirmed in different studies, albeit without further identification of which domain facilitates this interaction [9–11]. The spike protein of SARS-CoV-2 consists of several domains that contain a variety of functions. The S2 domain of the spike protein contains the viral fusion machinery. The S1 domain is divided into an N-terminal domain (NTD) and a C-terminal domain, which is referred to as receptor-binding domain (RBD) and contains the receptor-binding site [12, 13]. The RBD gathered the most attention as it is prone to immune recognition, to inhibit receptor binding and thus infection. Continuous immune pressure resulted in the emergence of mutants escaping from neutralizing antibodies and/or with increased ACE2 binding affinity [14–21]. For the RBD a (sialo)glycan-dependent attachment mechanism is described, whereby sialic acid and heparan sulfate moieties function as an initial point of attachment [5, 8]. Several coronaviruses harbor a glycan-binding function in the NTD, and using this domain, coronaviruses can bind a wide variety of glycan structures from non-sialylated N-glycans to heavily modified sialic acids in glycolipids [22–24]. Recently, the NTD of SARS-CoV-2 has been shown to contain a sialic acid binding site [25, 26], and low-affinity binding has been demonstrated by using Saturation Transfer Difference (STD)-NMR methods. The NTD is also an important antigenic site [27, 28], indicating its critical role in the viral life cycle with an apparent function.

We wanted to examine whether antigenic drift in the NTD could result in improved sialic acid binding properties. We, therefore, created trimeric NTD of VoC spike proteins and analyzed their binding to Vero E6 cells and tissue slides from several species. Three prevalent variants of 501Y.V2 (Beta, B. 1.351) were circulating in South Africa originating from SARS-CoV-2 Wuhan with the D614G mutation, lineage 501Y.V2-1, 501Y.V2-2 and 501Y.V2-3 [29]. The early 501Y.V2-1 Beta subvariant gained a receptor-binding function using its NTD, which appeared to be sialic acid-dependent since sialidase treatment abrogated binding. This NTD-sialoglycan binding functionality was lost in the subvariant 501Y.V2-3, which is commonly referred to as B.1.351 and quickly became the dominant Beta variant.

Using glycan arrays, ESI-MS, STD-NMR methods, and a graphene-based sensor we here demonstrate that the SARS Beta subvariant 501Y.V2-1 NTD protein can engage 9-*O*-acetylated α2-8 linked disialic acids.

## Results

### The Beta NTD protein gain-of-function is sialic acid dependent

We started to test recombinant NTD proteins of different VoC as several other members of the Coronavirus family employ this protein domain to bind sialylated or non-sialylated glycans for attachment [7, 24, 30, 31]. Indeed, it is known that this domain contains a galectin fold and weak sialic acid-binding of SARS-CoV-2 NTD has been reported using STD-NMR methodologies [25, 26, 32]. However, using the fluorescent NTD trimers of the S1 spike domain, we were not able to detect binding of SARS-CoV-2 Wuhan NTD to cells and lung tissues as previously described [33]. Circulation of SARS-CoV-2 and continuous adaptation have led to the emergence of VoCs (Alpha, Beta, Delta and Omicron), with the more recent Omicron variant having an unusual number of mutational changes in the NTD (Fig. 1A). In particular, the NTDs were expressed with a trimerization domain and a C-terminal fluorescent mOrange2 reporter (Fig. 1B), while receptor binding was characterized on Vero E6 cells, commonly used to isolate/propagate SARS-CoV-2 viruses since these cells support viral receptor binding and replication [34]. For SARS-CoV-2 Wuhan, Alpha, Delta and Omicron NTD, no tangible signal was detected to Vero E6 cells by confocal imaging (Fig. 2A and Supplementary Fig. 1). On the other hand, the 501Y.V2-1 variant did bind efficiently to cells. This binding property appeared to be dependent on the presence of sialic acids since enzymatic treatment with sialidase abrogated cell binding (Fig. 2A). Binding of 501Y.V2-1 NTD was further compared to SARS-CoV-2 Wuhan and other VoCs on formalin fixed, paraffin embedded lung tissue slides from ferret, Syrian hamster and mouse (Supplementary Fig. 2). Similar to cell staining, efficient binding was observed using the 501Y.V2-1 NTD on the lung tissue slides of Syrian hamster and mouse, with minor staining to ferret lung tissue.

**Figure 1:**
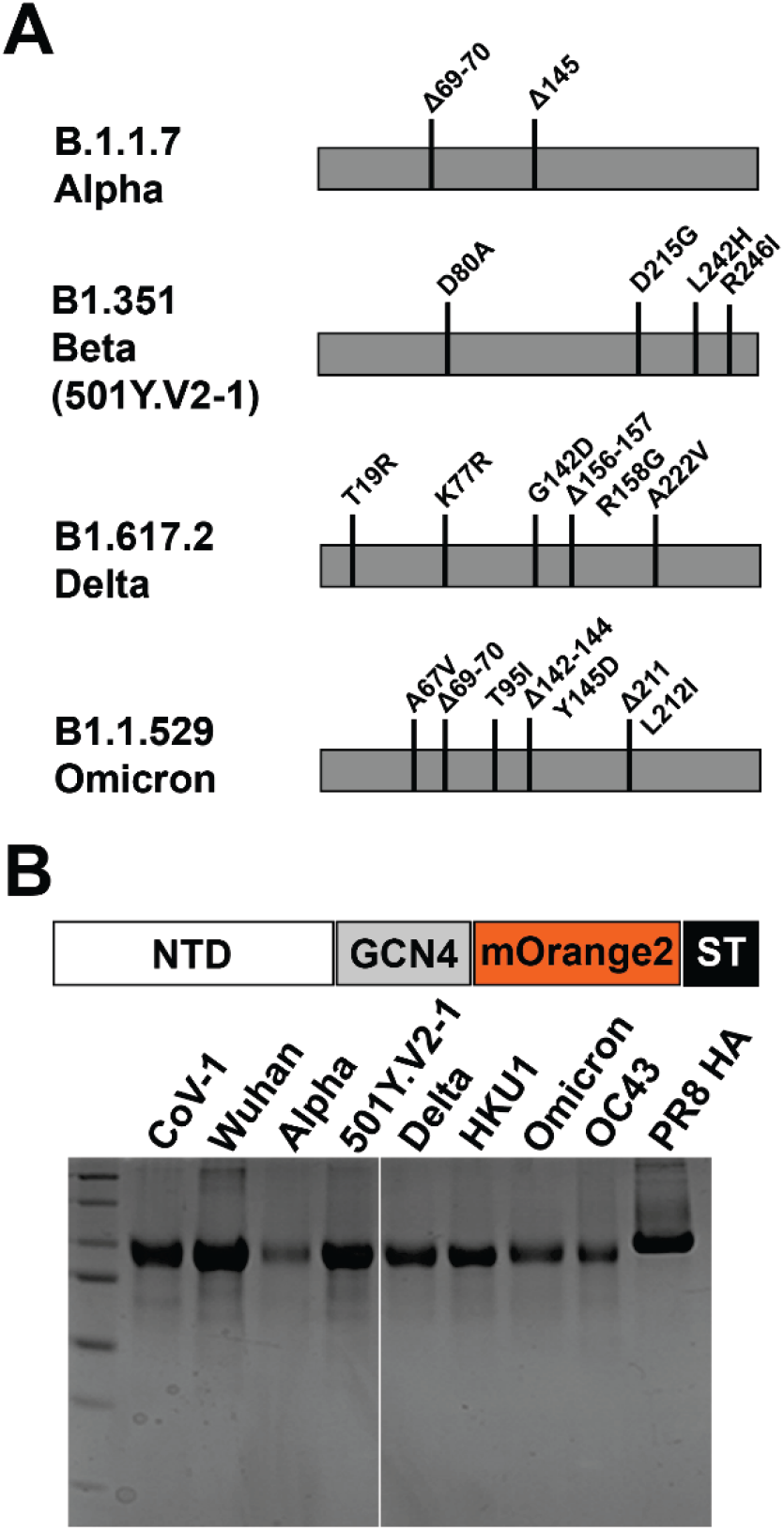
(A) Mutations in VoCs in relation to SARS-CoV-2 Wuhan. (B) NTDs expressed as trimeric proteins using a GCN4 trimerization domain, C-terminally fused to mOrange2 shown on coomassie gel.

**Figure 2:**
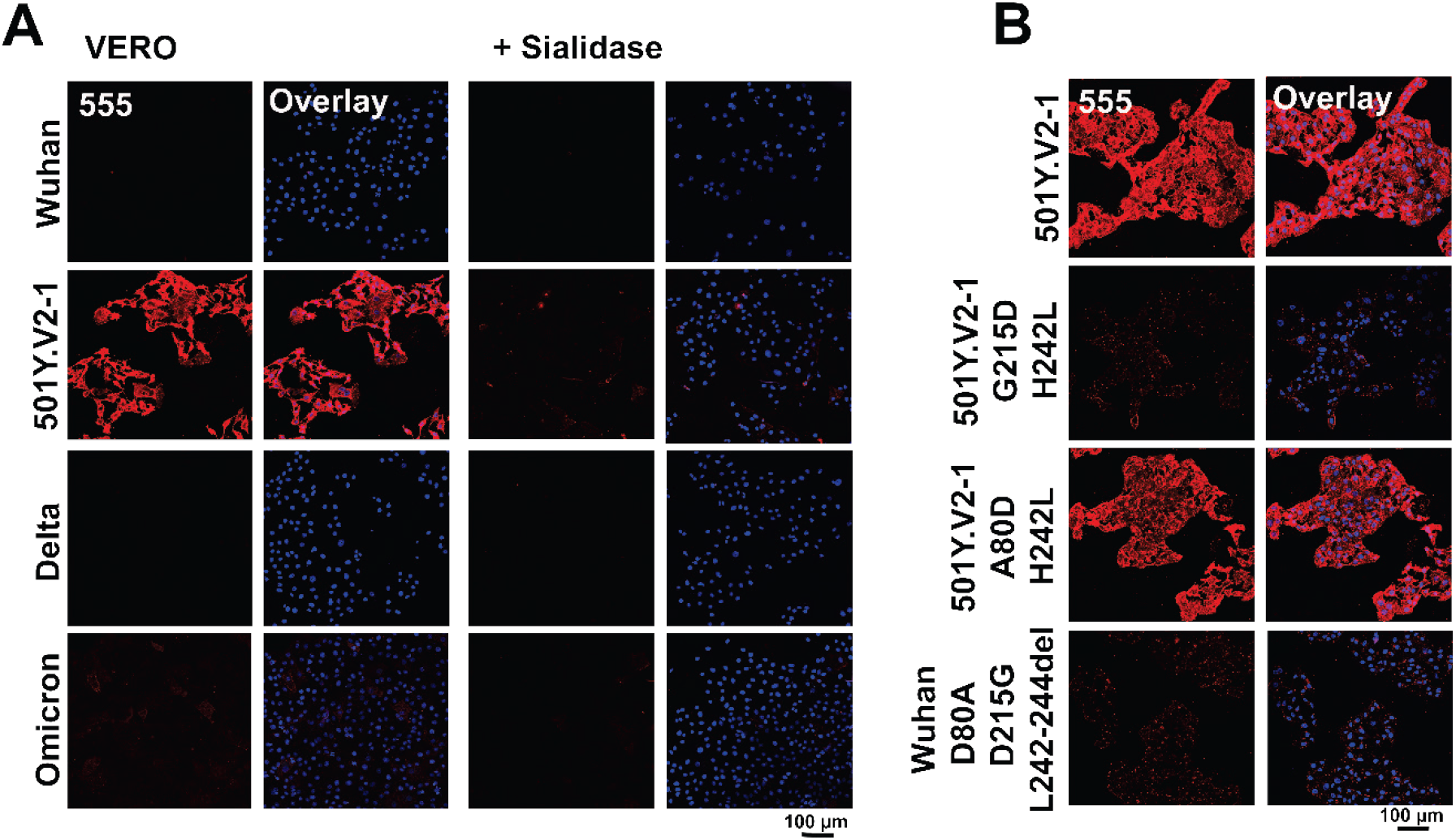
(A) SARS-CoV-2 VoC Beta (501Y.V2-1) trimeric NTD protein gaining sialic acid dependent cell binding, Beta NTD receptor binding abrogated with sialidase treatment, no binding observed for SARS-CoV-2 Wuhan, Delta and Omicron. (B) Mutations in 501Y.V2-1 NTD (G215D+H242L and A80D+H242L) NTD and in SARS-CoV-2 (D80A+D215G+L242-244del) delineate the importance of amino acid G215D and the amino acids at position 242-244 being important for receptor binding.

The Beta variant employed in these experiments retain the amino acids at position 242-244 with H242L (Fig. 1A); this non-dominant variant (501Y.V2-1) appeared to circulate initially in South Africa. Five amino acid mutations were identified in the 501Y.V2-1 variant: D80A, D215G, E484K, N501Y and A701V. Subsequent mutations were introduced into the 501Y.V2-1 variant S protein (L18F and K417N) which resulted in the emergence of 501Y.V2-2. Hereafter a deletion at 242-244 caused the dominant final variant to appear (501Y.V2-3) [35, 36]. The deletions at 242-244 result in the disruption of neutralizing monoclonal antibody binding, facilitating immune escape [37, 38]. These amino acids precede the N5-loop supersite (amino acids 246-260) to which potent antibodies are elicited. Additionally, amino acids at positions 242-244 appeared to be essential in stabilizing the cryptic SARS-CoV-2 NTD binding pocket and facilitate sialic acid interaction [26].

Further characterization was performed by generating mutants with different combinations in 501Y.V2-1 (G215D+H242L and A80D+H242L) and in SARS-CoV-2 (D80A+D215G+L242-244del) NTD to assess which amino acid-mutation/deletion is essential for sialic acid-dependent binding. Cell staining with 501Y.V2-1 G215D+H242L NTD resulted in a loss of cell binding, while staining with 501Y.V2-1 A80D+H242L NTD resulted in a similar receptor-binding capacity, as observed using 501Y.V2-1 NTD (Fig. 2B). For SARS-CoV-2 Wuhan NTD with mutations D80A, D215G and L242-244del no cell binding could be detected. Taken together, this data indicates that amino acid mutation D215G is essential and that deletion of amino acids at position 242-244 is detrimental for sialic acid-binding.

### SARS-CoV-2 501Y.V2-1 Variant of Concern exhibits an analogous binding specificity as other β-coronavirus N-terminal domains

Binding of 501Y.V2-1 NTD to Vero E6 cells and tissue slides and subsequent abrogation of binding by sialidase treatment instigated further characterization of receptor specificity. Glycan microarray technology was utilized to determine which sialylated structures can be bound. However, it must be noted that previous glycan microarrays [8, 39–43] were not able to fully characterize of Beta NTD-glycan interactions, as no binding was observed. Since spike proteins of human coronaviruses (OC43 and HKU1) bind to 9-*O*-acetylated sialic acids, characterization of receptor specificity of 501Y.V2-1 NTD was, therefore, assessed using a glycan microarray with a collection of *O*-acetylated sialoglycans to determine comparable specificity. Glycan microarrays without acetylated sialoglycans did not display any binding interactions. The SARS-CoV-2 Wuhan strain, Alpha, Delta and Omicron variant did not display any responsiveness to *O*-acetylated sialic acids, whereas for the 501Y.V2-1 variant NTD responsiveness was observed towards a α2-8 linked disialic acid structure containing 9-*O*-acetyl (**3g**) (Fig. 3 and Supplementary Fig. 3). This specificity was remarkably similar to HKU1. OC43 and Influenza D (OK/D) HEF displayed a much broader *O*-acetylated sialic acid receptor-binding capacity (Supplementary Fig 2).

**Figure 3.**
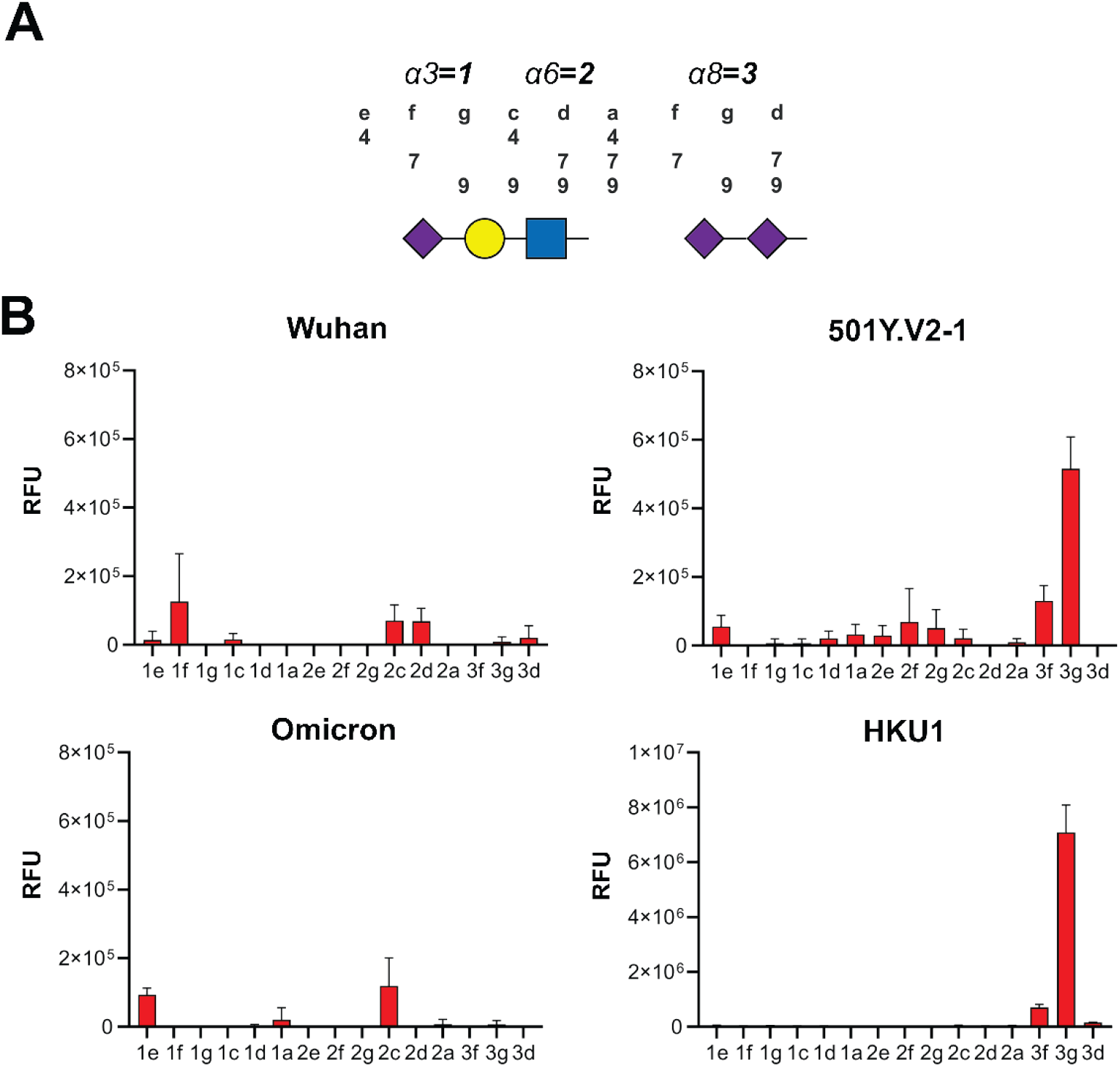
(A) 2-3, 2-6 and 2-8 linked sialic acid structures with 4, 7, 8 or 9 O-acetylation. (B) 501Y. V2-1 NTD gains binding to 9-O-acetylated 2-8 linked disialic acid similarly to HKU1 NTD with no binding being observed for SARS-CoV-2 Wuhan and Omicron VoC.

### CaR-ESI-MS and electrochemical sensor screening of 501Y.V2-1 NTD to assess receptor specificity towards 9-O acetylated disialosides

Further characterization of 9-*O*-acetylated disialosides responsiveness was performed using catch-and-release electrospray ionization mass spectrometry (CaR-ESI-MS) experiments to verify the specificity of 501Y.V2-1 NTD. CaR-ESI-MS is a semi-quantitative highly sensitive screening technique that allows for label-free detection of weak glycan-lectin interactions. Identification of ligands is achieved after their release (as ions) from glycan-lectin complexes following collision activation in the gas phase [8, 44, 45]. This approach was used to screen the specificity of SARS-CoV-2 Wuhan, 501Y.V2-1, Delta, Omicron and HKU1 NTDs against 9-*O*-acetyl α2-8 linked disialic acid or 9-*O*-acetyl α2-3-sialyllactosamine, which did not display any responsiveness on the glycan microarray. For all screening experiments, aqueous solutions composed of the volatile ammonium acetate salt (200 mM, pH 7.4, 25 °C), NTD (10 μM) and 9-*O*-acetyl α2-8 linked disialic acid or 9-*O*-acetyl α2-3-sialyllactosamine (10 nM of each glycan) were used. Representative CaR-ESI mass spectra acquired in negative mode are shown in Fig. 4. NTD-ligand complex ions with m/z values in the range of 5,000-7,000 were isolated for HCD. Over a range of collision energies (120-160 V), 9-*O*-acetyl α2-8 linked disialic acid was released, intact, as deprotonated (m/z 1065.45) or sodium adduct ions (m/z 1087.43) from the 501Y.V2-1 (highlighted region), Omicron and HKU1 NTD-ligand complexes. Notably, CaR-ESI-MS screening of five NTDs against 9-*O*-acetyl α2-3-sialyllactosamine produced no detectable released ligand signal, thus providing a blank experiment.

**Figure 4.**
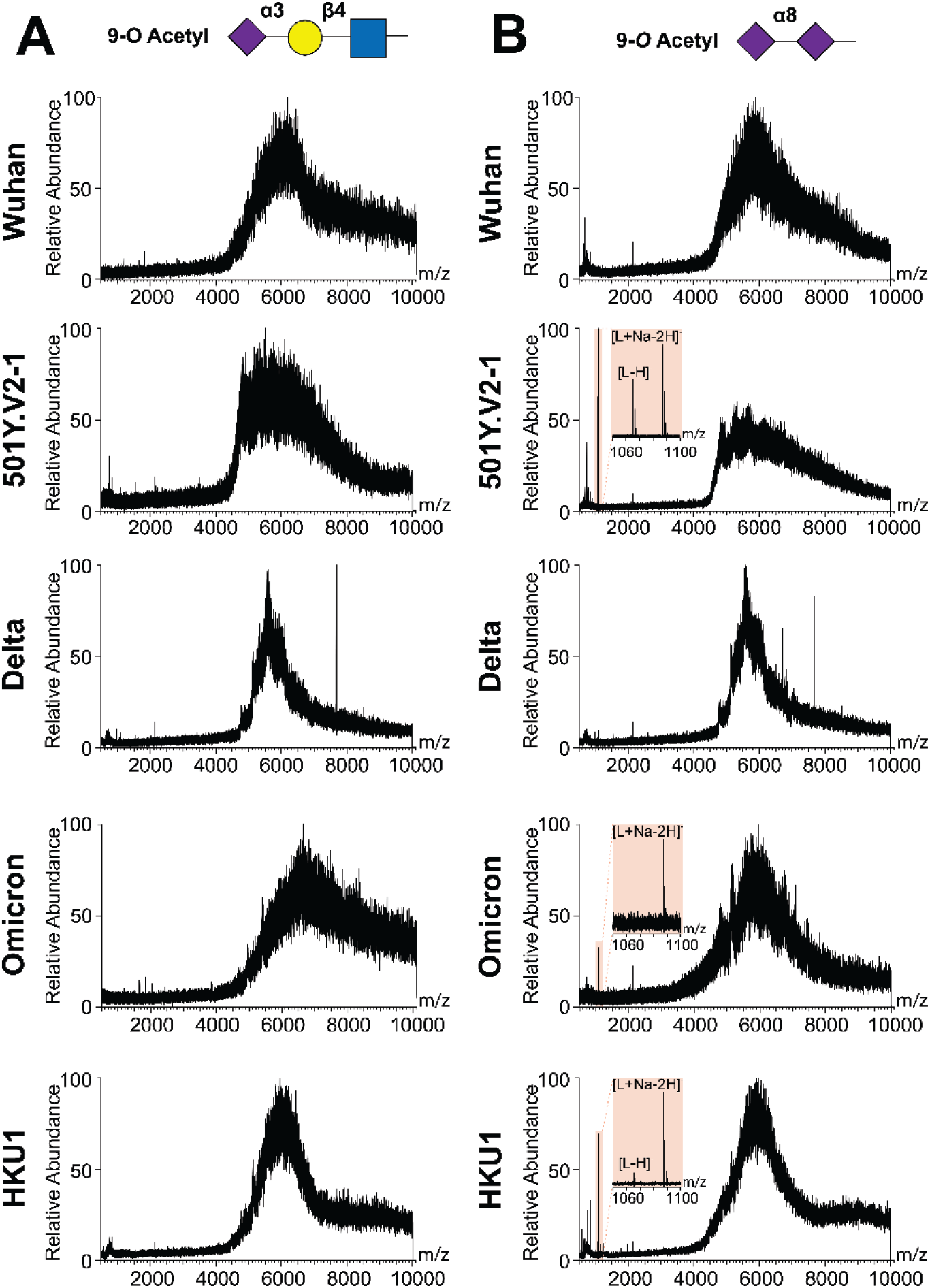
ESI-MS analysis confirms 9-O-acetylated sialic acid binding. CaR-ESI-MS screening results obtained for aqueous ammonium acetate solutions (200 mM, pH 7.4, 25 °C). SARS-CoV-2 Wuhan, 501.V2-1, Delta, Omicron and HKU1 NTDs in combination with 10 nM of 9-O-Ac 3-sialyllactosamine (A) or 9-O-Ac α2-8 disialic acid (B). Ions with m/z of 5,000–7,000 were subjected to HCD using a collision energy of 160 V. Screening with 9-O-acetyl α2-3-sialyllactosamine using five NTDs did not result in glycan release, whilst for 501Y.V2-1, Omicron and HKU1 NTD glycan release was observed with 9-O-acetyl α2-8 linked disialic acid (highlighted region).

Further analysis was performed with a screen-printed electrode sensor using differential pulse voltammetry. This technique characterizes the oxidation in peak current as a function of the ferro/ferricyanide redox reaction that is influenced by the barriers created on the electrode surface (i.e. ligand to protein binding) by obstructing diffusion of [Fe(CN)6]^3-/4-^[46, 47]. For the surface coated SARS-CoV-2 Wuhan, Delta, and Omicron NTD no signal was detected when using non-acetylated α2-8 disialic acid and 9-*O*-acetylated α2-8 disialic acid structures (Fig. 5). Delta NTD occasionally displayed minimal background signal. The strongest binding signals were detected for HKU1 NTD to 9-*O*-acetyl α2-8 linked disialic acid followed by 501Y.V2-1 NTD, thus further verifying the glycan-lectin interaction of 501Y.V2-1 and HKU1 NTD to the 9-*O*-acetylated α2-8 linked disialic acid in a concentration-dependent manner. Binding of 501Y.V2-1 and HKU1 NTD was not detected towards non-acetylated α2-8 disialic acid (Fig. 5).

**Figure 5.**
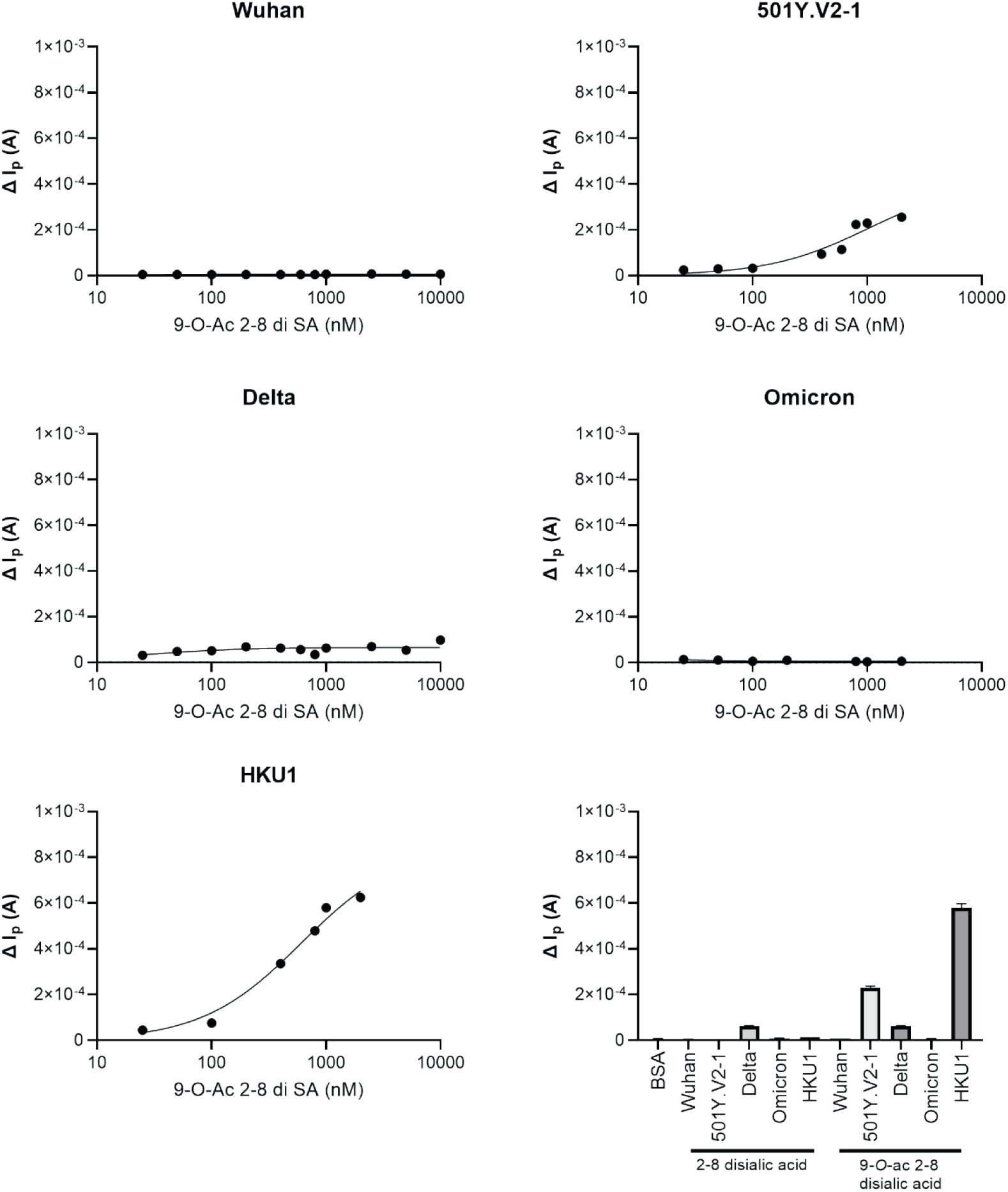
Graphene-based biosensor verifies 501Y. V2-1 NTD and 9-O-acetylated 2-8 linked disialic acid interaction. Screening of SARS-CoV-2 Wuhan, 501Y.V2-1, Delta, Omicron and HKU1 NTD using graphene-based biosensor against 9-O-acetylated α2-8 linked disialic acid in a concentration-dependent manner, with positive correlation being observed for 501Y. V2-1 and HKU1 NTD. Use of non-acetylated α2-8 linked disialic acid did not result in glycan-protein interaction for SARS-CoV-2 Wuhan, 501Y. V2-1, Delta, Omicron and HKU1 NTD, whilst signal is detected for Beta and HKU1 NTD using 9-O-acetylated α2-8 linked disialic acid.

### Identifying the binding epitope of 501Y. V2-1 NTD on the 9-O-acetylated ligand

Glycan microarray, CaR-ESI-MS and electrochemical analysis elucidated to which structure 501Y.V2-1 NTD binds to. Next, STD-NMR was employed, which allows for the identification of the binding epitopes within the ligand to a target receptor [48–50]. In STD-NMR, some protons within the protein are selectively irradiated with low power radiofrequency. Under spin diffusion conditions the magnetization is quickly transferred to all the protons of the receptor, which results in efficient protein saturation. If binding occurs, magnetization is transferred from the protein to the ligand protons. Importantly, not all protons of the ligand receive the same amount of saturation. Protons that are in closer proximity to the protein receive the strongest saturation, whilst the more distant will receive low saturation or none. Therefore, the resulting STD-NMR spectrum, which only contains the ligand protons that are affected by the protein saturation, not only detects binding or non-binding, but also informs about which part of the ligand is in closer contact with the protein. Thus, ^1^H STD-NMR experiments were performed to determine whether the SARS-CoV-2 Wuhan, 501Y.V2-1, Omicron and HKU1 NTDs bind the 9-*O*-acetyl α2-8 linked disialic acid and to define the corresponding ligand epitope (Fig. 6 and 7). The results from this analysis showed that under these experimental conditions (in solution, using a relatively high ligand 0.8 mM concentration) SARS-CoV-2 Wuhan, 501Y.V2-1 and HKU1 NTD indeed recognize 9-*O*-acetyl α2-8 linked disialic acid as a ligand. For SARS-CoV-2 Wuhan and 501Y.V2-1, the main ligand epitope is the terminal sialic acid with the strongest ^1^H STD-NMR signals arising from the 9-*O*-acetylated glycerol chain, indicating that

**Figure 6.**
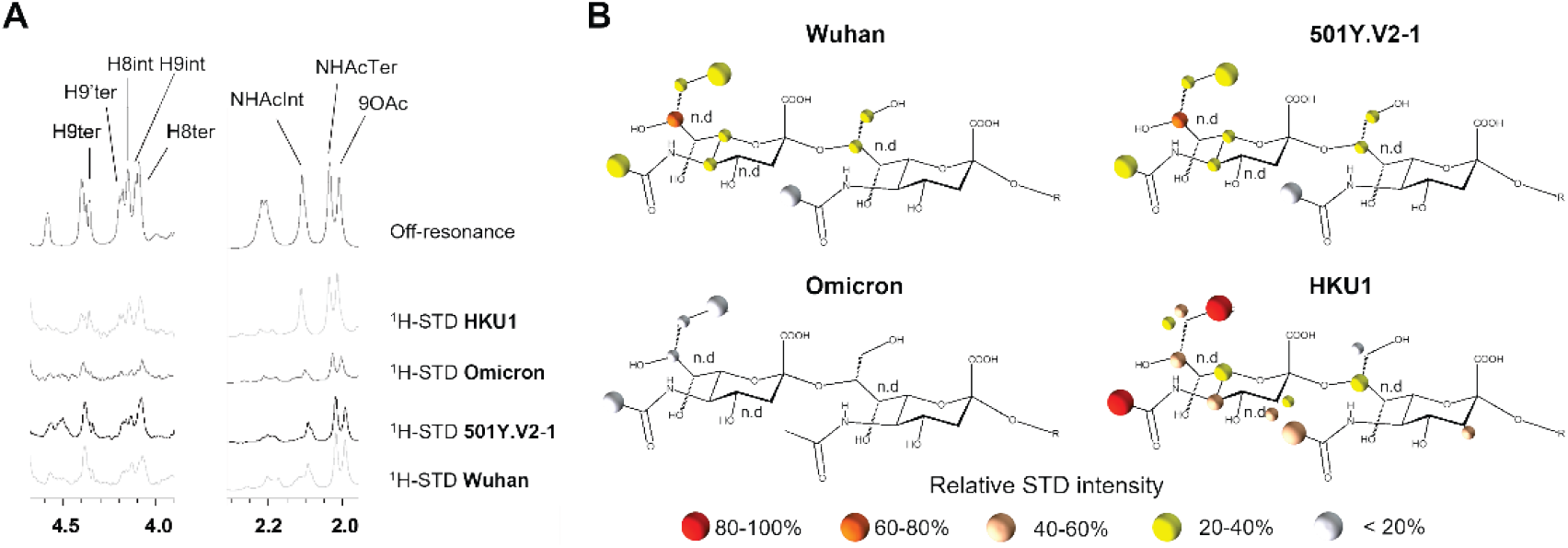
^1^H STD NMR experiments for the interaction of the spike proteins of SARS-CoV-2 variants and HKU1 with the 9-O-acetyl α2-8 linked disialic acid. (A) Selected area of ^1^H STD-NMR spectra with aliphatic protein irradiation. (B) Ligand epitope mapping presented as relative STD intensities.

**Figure 7.**
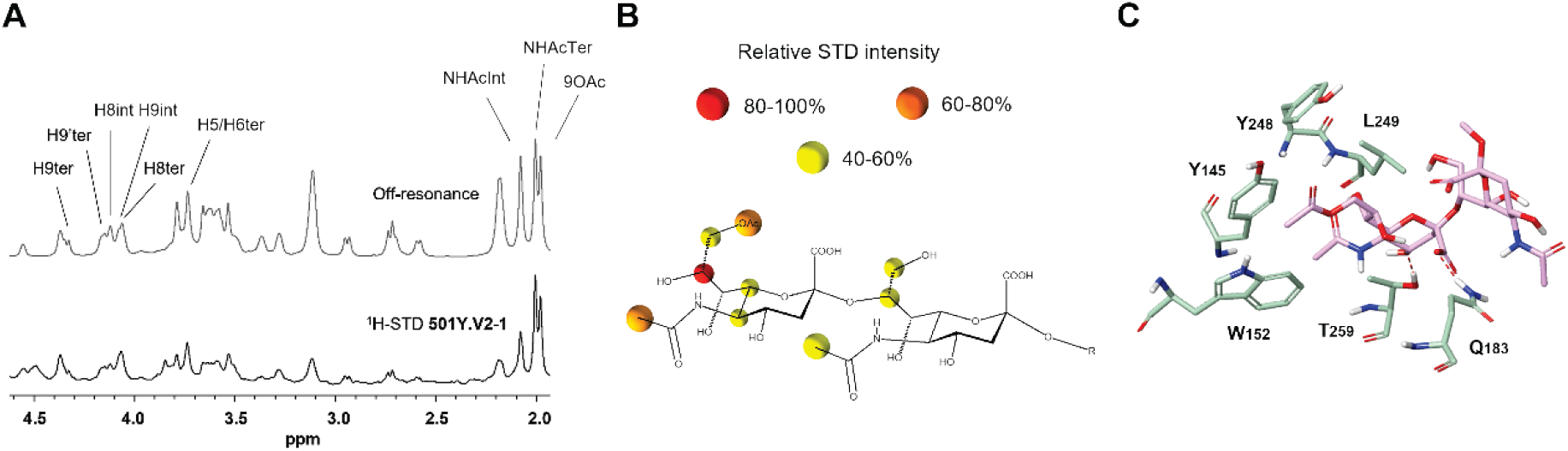
Interaction of the spike proteins of SARS-CoV-2 501Y.V2-1 with 9-O-acetyl α2-8 linked disialic acid. (A)^1^H STD NMR experiment. (B)Ligand epitope mapping presented as relative STD intensities. (C) Molecular model as derived by docking of the ligand into the sialic acid binding site. PDB code 7QUR^[23]^.

this fragment is tightly interacting with these NTD proteins. Additional STD-NMR signals were detected from the methyl groups of the acetamide group at C-5 of both sialic acid moieties. Medium-weak STD-NMR signals were also observed from the H-5 and H-6 of the terminal sialic acid, while the protons at the C-3 provide STD-NMR signals neither in the terminal nor in the reducing end sialic acids. To complement these STD-NMR experimental derived results, a putative complex of the NTD bound to 9-*O*-acetyl α2-8 linked disialic acid was built by using molecular modeling tools (Fig. 7C). According to the generated model, the carboxylate of the terminal sialic acid establishes a hydrogen bond interaction with the Q183 side chain, the acetyl moiety at the C-9 fits into a protein hydrophobic pocket defined by residues W152 and Y145, while the acetamide group at the C-5 is accommodated into a second pocket flanked by L249 and T259. Finally, the acetamide of the sialic acid at the reducing-end also faces the protein surface, although no specific intermolecular interactions were found. The same analysis was performed for the Omicron variant. Minimal ^1^H STD-NMR signals were detected for this variant in comparison to SARS-CoV-2 Wuhan, suggesting that the Omicron NTD has lost its ability to efficiently recognize the 9-*O*-acetyl α2-8 linked disialic acid molecule as a ligand (Fig. 6B). Finally, the NTD of HCoV-HKU1 was characterized, consistently, the ^1^H STD-NMR spectrum of the corresponding complex with the 9-*O*-acetyl α2-8 linked disialic acid displayed the strongest STD intensities with the main ligand epitope involving the acetyl moieties of the terminal sialic acid. Medium-strong STD-NMR signals were detected for the glycerol chain and for H-5 and H-3 axial of the terminal sialoside, as well as for the acetamide moiety and the H-3 equatorial of the reducing end sialoside. Medium-weak STD signals were also recorded for one of the H9, H-6 and H-3 equatorial of the terminal sialoside and for the H-8 of the internal residue. In all cases, while some of the STD-NMR signals could not be properly quantified due to ^1^H-NMR signal overlap, the NTD-9-*O*-acetyl α2-8 linked disialic acid interaction was unambiguously proven.

## Discussion

SARS-CoV-2 spike displays differential abilities to bind to glycans [5, 8]; for SARS-CoV-2 NTD only a weakly binding glycan site is observed when using highly sensitive methods [25, 26]. In this report, we have characterized, with biochemical assays, the glycan binding properties of NTDs of SARS-CoV-2 VoCs. The 501Y.V2-1 NTD is the first VoC that displays a clear receptor-binding capacity; the D215G mutation and conservation of amino acids at position 242-244 appeared to be essential for sialic acid-dependent binding. Glycan microarray, CaR-ESI-MS, STD-NMR and electrochemical analysis validated 9-*O*-acetylated α2-8 disialic acid binding specificity of the 501Y.V2-1 NTD. The 501Y.V2-1 variant shows strong interaction with the 9-*O*-acetyl and NHAc-5 arm NTD of the ligand, similarly to HKU1, indicating a convergent evolution of coronaviruses towards host adaptation and 9-*O*-acetylated sialoside recognition using their NTD [7, 11, 23, 25, 26, 51–54]. Omicron NTD interaction with 9-*O*-acetylated 2-8 disialic acid was minimally detected with CaR-ESI-MS, weak binding was observed with STD-NMR and confocal imaging, whereas binding was not observed with a glycan array and a graphene-based biosensor. Low-affinity interaction with acetylated structures may be possible for other VoCs, however, this could not be validated.

Initial virus-receptor interaction may vary in affinity and avidity, whereby engagement first occurs with a low-affinity/high-avidity interaction (glycan binding) and then potentially followed by a stronger secondary receptor interaction (protein binding) [55]. Many viruses utilize reversible low-affinity interaction through binding with (sialylated) glycans to mediate viral surfing [3, 56, 57]. Coronaviruses utilize their NTD for sialic acid binding and promote infection [7, 24, 30, 31, 54, 58]. So far, only MERS-CoV has been identified as using a two-step attachment mechanism that contributes toward tropism, by utilizing its S1-NTD to interact with α2-3 sialic acid structures (viral surfing) [59]. Even though a difference in the sequence is observed among HCoV-HKU1, MERS-CoV and SARS-CoV-2, these proteins share high protein folding similarity and structural overlap whereby the sialic acid-binding site appears to be conserved [7, 60–62]. In-silico structural and molecular studies elaborate that the flat and non-sunken NTD surface of SARS-CoV-2 should allow for sialic acid interactions, enabling viral surfing and dual-receptor functionality of the S1 spike [63–65]. S1 spike interaction with sialic acids has been further shown by four independent studies, however, the contribution of S1-NTD towards sialoside recognition and dual-receptor functionality has not been verified [8, 10, 11, 66].

STD-NMR experiments [26] have identified a clear “end-on” interaction of S1-NTD with α2-3- or α2-6-linked sialic acids, describing a cryptic sialic acid binding domain. A cryo-EM-derived map of the S1-NTD has proposed H69, Y145 and S247 as the sialic acid-interacting amino acids [26]. The amino acid at position Y145 interacts with the glycerol moiety (C7-C9) of the terminal sialic acid with a hydrogen bond and the S247 amino acid engaging the NHAc-5 arm, which is also supported by computational modeling described in our report. Numerous mutations in the sialoside binding site of VoCs appear to remove essential interacting residues (H69, V70, Y145: Alpha and Omicron) or perturb the β-sheet structure that forms the binding pocket (L242-244del: Beta). Interestingly, when using α2-3-, α2-6- and α2-8-linked *O*-acetylated sialic acids in our glycan-array, binding was only observed for 9-*O*-acetyl α2-8-linked sialic acids when using 501Y.V2-1 NTD. Previously inhibition of S1-spike binding was attained with Neu5Ac monosaccharides [26, 66], α2-3 linked [26], (multivalent) α2-6 linked [10, 26] and (multivalent) 9-*O*-acetylated sialic acid [11], which raises questions regarding the specificity of SARS-CoV-2 NTD towards these structures and whether sufficient receptor binding can be mediated to support a dual-receptor function. Thus, the current reported knowledge indicates that sialic acid binding and multivalency is crucial in infection and tropism whilst the exact glycan partner can vary between coronavirus strains [67].

The lectin function for 501Y.V2-1 VoC is a clear convergent evolutionary path of adaptation to 9-*O*-acetylated sialoside recognition by the NTD, similarly to β-CoV HKU1 and OC43 [7, 11, 23, 25, 26, 51–54]. However, ablation in later Beta/501Y.V2 variants induced by the L242-244deletion [35, 36] and in other VoC (Alpha, Delta and Omicron) raises the question of why this increased binding capacity towards sialic acids is lost. Influenza, HKU1 and OC43 possess a receptor-destroying enzyme (neuraminidase and hemagglutinin-esterase) that results in the “release” of virions from host cells [68]. OC43 without a hemagglutinin-esterase (HE) activity and influenza viruses without neuraminidase activity have poor viral dissemination, by preventing release from infected host cells [69, 70]. This may explain why in the dominant Beta/501Y.V2-3 variant enhanced sialic acid binding was abrogated. As SARS-CoV-2 does not possess HE on its viral membrane [71], a balance is needed between the ability to bind host cells for initial infection (tropism) and dissemination in the host (viral release). Similar to SARS-CoV-2, the MERS-CoV strain does not possess HE activity, although a dual-receptor functionality has been observed through α2-3-linked sialic acid interaction of the NTD. Interestingly, this interaction of MERS-CoV NTD with sialic acids appears to be extremely weak compared to HKU1 and OC43 NTD. Multimerization with a nanoparticle is required for MERS-NTD to observe sialic acid binding [59] whilst HKU1 and OC43 NTD have a strong affinity towards sialic acids [7, 24], suggesting that MERS-CoV retains a fine balance between sialic acid binding and subsequent release from infected cells using its NTD.

In summary, the observation that 501Y.V2-1 NTD has gained a lectin function supports the dual-receptor notion of S1 spike, similar to that of MERS-CoV, showing that SARS-CoV-2 might probe evolutionary space to allow for alternative or additional receptor binding. Subsequent ablation in VoC raises questions regarding the convergent evolutionary path of glycan binding proteins to recognize (acetylated) sialylated glycan structures and the fine balance it requires for infection and/or transmission.

## Material and methods

### SARS-COV-2 NTD expression plasmid generation

Recombinant SARS-CoV-2 spike protein N-terminal domain (GenBank: MN908947.3; AA319-541) was cloned using Gibson assembly from cDNAs encoding codon-optimized open reading frames of full-length SARS-CoV-2 spike [72], as previously described [33]. The pCD5 expression vector was adapted so that after the signal sequence, the SARS-COV-2 NTD-encoding cDNAs are cloned in frame with a GCN4 trimerization motif (KQIEDKIEEIESKQKKIENEIARIKK), a TEV cleavage site, fluorescent reporter open reading frame [73, 74], and the Strep-Tag II (WSHPQFEKGGGSGGGSWSHPQFEK); IBA, Germany).

### Protein expression and purification

pCD5-SARS-COV-2 NTD-+ GCN4 - Fluorescent probe expression vectors were transfected into HEK293T with polyethyleneimine I (PEI) in a 1:8 ratio (μg DNA:μg PEI) as previously described [75]. The transfection mix was replaced after 6 hours by 293 SFM II suspension medium (Invitrogen, 11686029, supplemented with glucose 2.0 gram/L, sodium bicarbonate 3.6 gram/L, primatone 3.0 gram/L (Kerry), 1% glutaMAX (Gibco), 1.5% DMSO and 2mM valproic acid). Culture supernatants were harvested 5 days post-transfection. The SARS-COV-2 NTD expression was analysed with SDS-PAGE followed by Western-blot on PVDF membrane (Biorad) using α-strep-tag mouse antibodies 1:3000 (IBA Life Sciences). Subsequently, SARS-COV-2 NTD proteins were purified with Sepharose Strep-Tactin beads (IBA Life Sciences) as previously described [75].

### Immunofluorescent cell staining

VERO E6, grown on coverslips were analyzed by immunofluorescent staining. Cells were fixed with 4% paraformaldehyde in PBS for 25 min at RT after which permeabilization was performed using 0.1% Triton in PBS. Subsequently, the coronavirus spike proteins were applied at 50 μg/ml for 1 h at RT. Primary StrepMAB-Classic-HRP (IBA) and secondary Alexa-fluor555 goat anti-mouse (Invitrogen) were applied sequentially with PBS washes in between. DAPI (Invitrogen) was used as nuclear staining. Samples were imaged on a Leica DMi8 confocal microscope equipped with a 10x HC PL Apo CS2 objective (NA. 0.40). Excitation was achieved with a Diode 405 or white light for excitation of Alexa555, a pulsed white laser (80MHz) was used at 549 nm, emissions were obtained in the range of 594-627 nm respectively. Laser powers were 10 - 20 % with a gain of a maximum of 200. LAS Application Suite X was used as well as ImageJ for the addition of the scale bars.

### Glycan array

Acetylated structures were printed on glass slides as previously described [24]. The glycan microarray utilized as described previously for NTD proteins [76]. Precomplexation was performed with NTD proteins using StrepMAB-Classic-HRP (IBA) and goat anti-mouse-Alexa555 antibodies in a 4:2:1 molar ratio respectively in 50 μl phosphate-buffered saline (PBS) with 0.1 % Tween 20. Samples were incubated on ice for 15 mins, followed by incubation on the array for 90 mins in a humidity chamber. Slides were rinsed with Tween 20, PBS and deionized water, followed by centrifugation and scanning as described previously [76]. Data processing was performed using six replicates, lowest and highest replicates were removed and subsequent mean and standard deviation were calculated using the remaining four replicates.

### ESI-MS

#### Proteins and glycans

Protein stock solutions were dialyzed against 200 mM aqueous ammonium acetate pH 7.4 using an Amicon 0.5 mL micro concentrator with a MW cutoff of 10 kDa (EMD Millipore, Billerica, MA) and stored at 4 °C until needed. The concentration of each protein stock solution was estimated by UV absorption at 280 nM.

Stock solutions of each glycan were prepared by dissolving a known mass in 100 mM ammonium bicarbonate (pH 7.4) with ultrafiltered water (Milli-Q Millipore, MA) to achieve final concentration of ~ 1 mM. All stock solutions were stored at −20 °C until used.

#### Mass spectrometry

The CaR-ESI-MS experiments were performed in negative mode using a Q Exactive Ultra-High Mass Range Orbitrap mass spectrometer (Thermo Fisher Scientific). The mass spectrometer was equipped with a modified nanoflow ESI (nanoESI) source. NanoESI tips with an outer diameter (o.d.) of ~ 5 μm were pulled from borosilicate glass (1.0-mm o.d., 0.78-mm inner diameter) with a P-1000 micropipette puller (Sutter Instruments). A platinum wire was inserted into the nanoESI tip, making contact with the sample solution. A voltage of approximately –1 kV was applied to the platinum wire.

The capillary temperature was 150 °C, and the S-lens RF level was 100; an automatic gain control target of 1 × 10^6^ and maximum injection time of 200 ms were used. The resolving power was set to 25,000. HCD spectra were acquired using collision energies ranging from 10 V to 300 V. Argon was used for collision-induced dissociation (CID) at a Trap ion guide pressure of 1.42 × 10^-2^ mbar. Data acquisition and preprocessing was performed using Xcalibur version 4.1.

### Graphene-based biosensor

#### Fabrication of the electrochemical graphene-based biosensor

A 10 μl mixture of 1 μM MUA and 10 μM DTT was dropped onto the AuNPs/G /GCE surface of the electrode and placed in a refrigerator (4 °C) for 14 h to obtain the MUA/AuNPs/GGCE. The as-prepared electrode was activated in 100 μl of a freshly prepared solution containing 2 g/L EDC and 0.5 g/L NHS for 30 min to activate the carboxylic groups on MUA. Then, the activated electrode was immersed in 100 μl of lectin (10 μM) or spike protein (100 nM) solution for 1 h. The spike protein/MUA/AuNPs/G/GCE was immersed in 100 μl of 1% BSA for 30 min to inhibit nonspecific interactions and then the electrode was rinsed thoroughly to remove any adsorbed components. The spike protein/AuNPs/G/GCE was stored at 4 °C in PBS (pH 7.4).

#### Electrochemical measurements

Electrochemical measurements were performed in 100 μl of analyte which includes 10 mM PBS containing 25 mM [Fe(CN)6]3-/4- and 0.2 M KCl. Cyclic voltammetry (CV) was used to monitor the fabrication process of the biosensor. All the CV voltammograms were recorded from −0.2 V to 0.8 V (vs. Ag/AgCl) at a scan rate of 0.05 V/s. Differential pulse voltammetry (DPV) was used as the validation method. All DPV voltammograms were recorded −0.2 V to 0.5 V (vs. Ag/AgCl) at a modulation time 0.05 s, modulation amplitude 0.1 V and interval time 0.5 s. All electrochemical experiments were performed at room temperature (25 ± 1 °C). Glycan ligands were added to reach a final concentration of 10000 nM.

### STD-NMR

^1^H STD-NMR experiments were acquired on the Bruker 800 MHz spectrometer with a cryoprobe (Bruker, Billerica, MA, United States) at 298K. Proteins was buffer exchanged by 20 mM phosphate buffer (pD 7.5) containing 150 mM NaCl and 0.05% sodium azide in D2O. All the proteins were concentrated to a final concentration of 8 μM. The glycan ligands were then added to reach a final concentration of 800 μM, which lead to a protein/ligand ratio of 1:100. ^1^H STD-NMR spectra were acquired by using standard Bruker STD sequence (stddiffesgp.3) with 1152 scans in a matrix with 64K data points, in a spectral window of 12335.5 Hz centered at 2818 Hz. An excitation sculpting module with gradients was used to suppress the water proton signals, and a protein suppression spin lock filter of 40ms. Protein’s resonance selective saturation was reached by irradiating at −0.2 ppm (aliphatic residues) using a series of 40 Eburp2.1000-shaped 90° pulses (50 ms) for a total saturation time of 2 s, and a relaxation delay of 3 s. For the reference spectrum an irradiation frequency at 100 ppm was used. Control STD-NMR experiments were performed both for the only ligands and apo proteins using the same STD experimental setup. Spectra analysis determined the percentages of STD intensities as estimated by comparing the intensity of the signals in the STD spectrum with the signal intensities of the off-resonance spectrum. The STD intensities of the ligands in absence of the protein were subtracted. The STD-derived epitope maps are represented as the relative percentages of each ligand signal with respect to the highest one. Resonances are labelled as n.d (not determined) when the ^1^H-NMR signal degeneration hampers rigorous quantitative analysis.

## Acknowledgements

R.P.dV is a recipient of an ERC Starting Grant from the European Commission (802780) and a Beijerinck Premium of the Royal Dutch Academy of Sciences. R.W.S. acknowledges support from the Netherlands Organization for Scientific Research (NWO) through a Vici grant and from the Bill & Melinda Gates Foundation grants INV-002022 and INV-008818. G.J.B is supported by the National Institutes of Health (P41GM103390 and R01HL151617) and by the Netherlands Organization for Scientific Research (NWO TOPPUNT 718.015.003). dr. M.A. Wolfert (Utrecht University) developed, printed and validated the glycan microarray. J.JB thanks the Agencia Estatal de Investigación (Spain) for Grant RTI2018-094751-B-C21, and CIBERES, an initiative of Instituto de Salud Carlos III (ISCIII), Madrid, Spain.

