## supplementary figures for "The SARS-CoV-2 spike N-terminal domain engages 9-*O*-acetylated α2-8-linked sialic acids"

Supplementary Fig 1: Confocal Vero e6 staining

*Supplementary fig 1: Confocal imaging of Alpha VoC NTD did not display any cell-binding properties.*

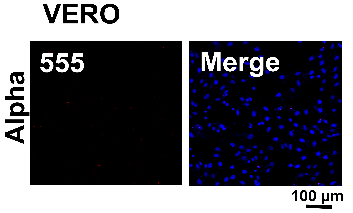

Supplementary Fig 2: Tissue staining

*Supplementary fig 2: Confocal imaging of SARS-CoV-2, Alpha, 501.V2-1, Delta and Omicron NTDs on ferret, Syrian hamster and mouse tissue slides. Binding of 501.V2-1 NTD was observed for all stained animal slides. For Wuhan minimal fluorescence signal was observed, similarly to the antibody control, explicit binding could not be verified.*

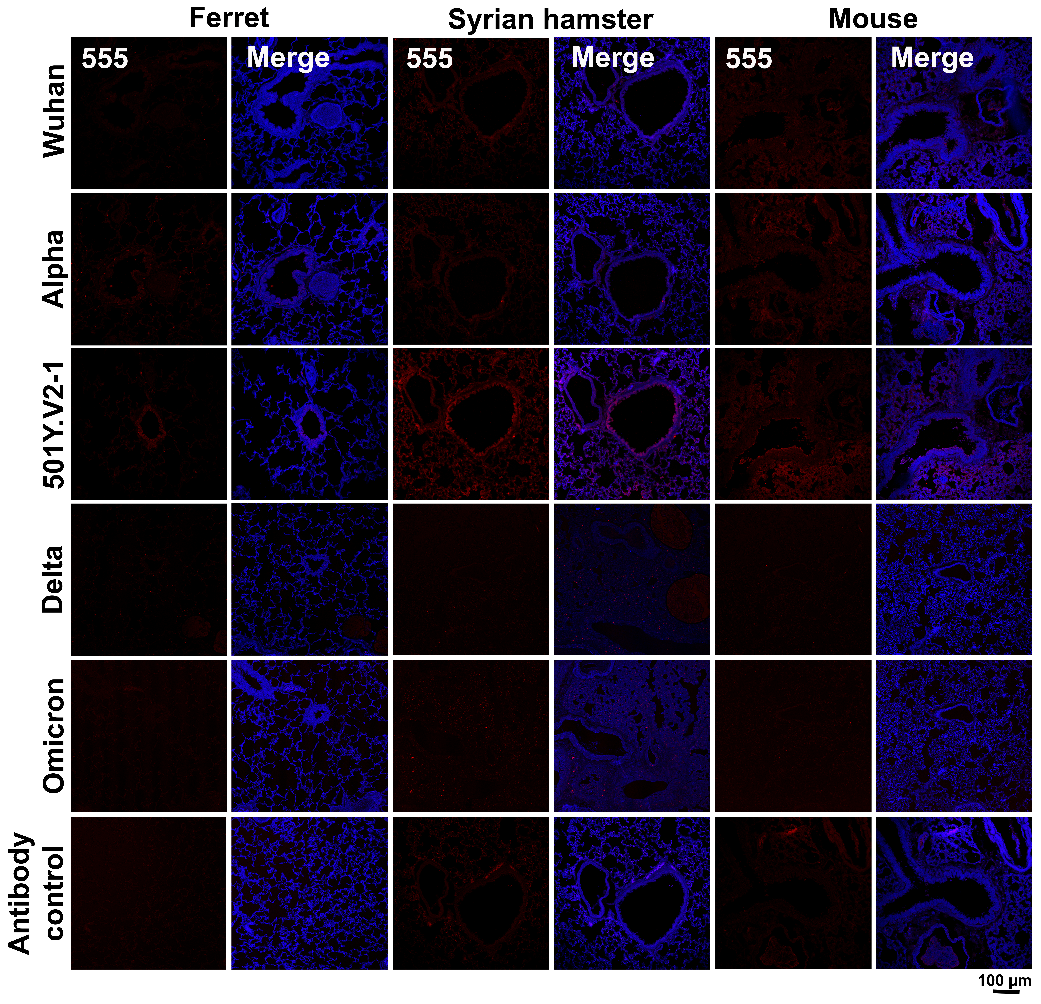

Supplementary Fig 3: Glycan array

*Supplementary fig 3: (A) O-Acetylated glycan structures. (B) O-acetyl glycan array analysis did not elucidate binding capacity to acetylated structures using Alpha or Delta VoC NTDs, with only background signal being observed. OC43 and OK/D show specificity towards acetylated structures.*

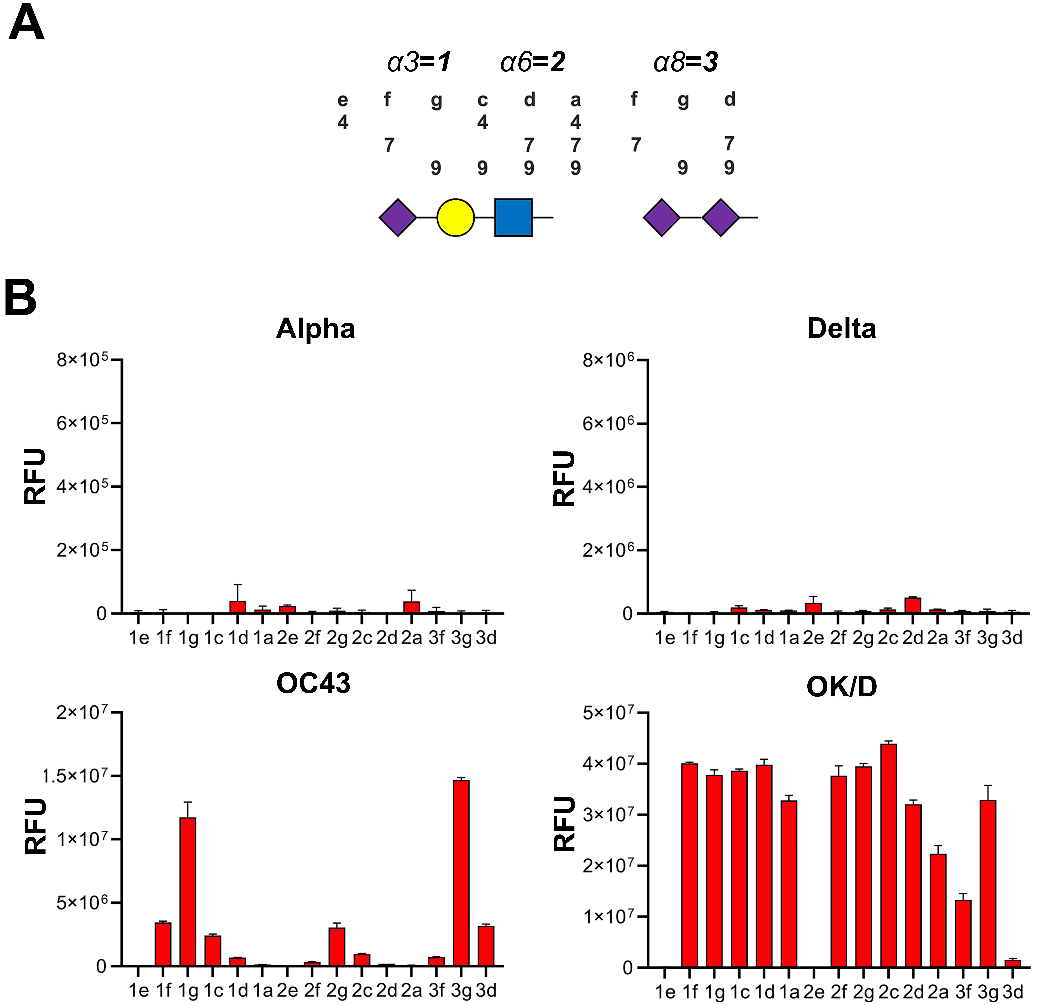
